# *fhl2b* expression ameliorates muscular dystrophy

**DOI:** 10.1101/2023.10.26.563896

**Authors:** Nils Dennhag, Abraha Kahsay, Itzel Nissen, Maria Chermenina, Hanna Nord, Jiao Liu, Anders Arner, Jing-Xia Liu, Ludvig J Backman, Silvia Remeseiro, Jonas von Hofsten, Fatima Pedrosa Domellöf

## Abstract

In muscular dystrophies, muscle fibers loose integrity and die, leading to significant suffering and a shorter life. Strikingly, the extraocular muscles (EOMs), controlling eye movements, are spared and function well despite the disease progression. Although EOMs have been shown to have important differences compared to body musculature the mechanisms underlying this inherent resistance to muscle dystrophies remain largely unknown. Here, we demonstrate important differences in gene expression as a response to muscle dystrophies between the EOMs and trunk muscle in zebrafish via transcriptomic profiling. We show that the LIM-protein Fhl2 is upregulated in response to knockout of *desmin*, *plectin* and *obscurin*, intermediate filament proteins whose knockout causes different muscle dystrophies, and contributes to disease protection of the EOMs. Moreover, we show that ectopic expression of *fhl2b* can partially rescue the muscle phenotype in the zebrafish Duchenne muscular dystrophy model *sapje*, significantly improving their survival rate. Therefore, *fhl2* is a protective agent and a candidate target gene for therapy of muscle dystrophies.

## Introduction

Muscular dystrophies, caused by mutations in more than 40 genes^1^ share similar features including muscle fiber disruption leading to muscle weakness, loss of ambulation and often premature death predominantly due to respiratory failure. Although muscular dystrophies affect 1:4-5000 births worldwide^2^ and induce pronounced suffering, there is currently no cure and treatment options are sparse. Thus, there is a need for new effective treatment options to prolong and improve the quality of life of patients suffering from these detrimental diseases.

The most common form of muscular dystrophy is Duchenne muscular dystrophy (DMD). In DMD, the giant protein dystrophin is lost or truncated^3,4^. Dystrophin is a crucial member of the dystrophin-glycoprotein-complex (DGC). The DGC links the extracellular matrix, across the cell membrane, to F-actin in the cytoskeleton within the myofibrils, which is fundamental for myofiber integrity^5^. The DCG contributes to multiple functions in the myofiber such as force transmission across the sarcolemma, but also acts as a signaling hub^6-8^. Dystrophin is therefore a key element for myofiber integrity. The most frequently used zebrafish DMD model is the *sapje* line^9,10^. *Sapje* zebrafish carry an A-to-T transversion in exon 4 of the dystrophin gene, resulting in a premature stop codon^9^. The lack of Dystrophin in zebrafish subsequently leads to detachment of trunk myofibers from the myosepta and failure of the contractile apparatus which ultimately results in premature death of up to 50% of zebrafish larvae at the age of 5 days post fertilization (dpf)^11^. Hence, the *sapje* zebrafish are severely affected by the lack of Dystrophin, but generally mimic the human DMD condition and constitute a good model for experimental treatment studies.

The EOMs have shown an innate resistance towards muscular dystrophies^12-15^. Even though EOMs share attributes with other striated skeletal muscles they differ in terms of gene expression and protein content^16,17^ as well as neuromuscular and myotendinous junction composition^18,19^. We have previously shown that zebrafish EOMs are a good model to study the cytoskeleton as well as myofiber and neuromuscular junction cytoarchitecture^20^. Recently, the muscle specific intermediate filament protein Desmin was shown to be naturally lacking in a subset of EOM myofibers in human and zebrafish^20,21^. Additionally, other intermediate filament proteins such as Nestin and Keratin-19 show a complex pattern in human EOM myotendinous junctions^19^. Altogether these findings suggest that the cytoskeletal composition of the EOMs differs from that of skeletal muscles in other parts of the body.

Desmin is the most abundant cytoskeletal protein in skeletal muscle fibers^22^ and, besides its interaction with the DCG, Desmin has been suggested to contribute to gene regulation^23^ and also as a mechanosensor, utilizing mechanical stretch to trigger intracellular signaling^24^. Additionally, Desmin anchors myonuclei and mitochondria to the sarcolemma and myofibrils^25^, reviewed in^26,27^. Despite this important role in the myofiber, studies of *Des* knockout mice have shown that the EOMs are relatively unaffected^28^. Patients with desminopathy have near normal life expectancy, with minor complications compared to DMD patients^29,30^. Desmin mutant animal models therefore offer a route to study the EOMs in a dystrophic background over extended periods of time. We hypothesized that the transcriptome of the EOMs in a dystrophic setting would provide information regarding their innate resistance towards muscular dystrophies, and that this could be utilized to rescue other dystrophic skeletal muscle.

In the current study, we used a *desmin* knockout zebrafish line to identify a subset of genes specifically upregulated in EOMs, via transcriptome analysis. We further investigated one of these genes, *fhl2b*, and demonstrate that *fhl2b* has an important role in the inherent resistance of EOMs towards muscular dystrophies. Additionally, we show that *fhl2b* significantly improves survival, muscle integrity and function of *sapje* zebrafish making it a novel target in the treatment of muscular dystrophies.

## Results

### A zebrafish model of desminopathy displays skeletal muscle defects

To study the EOMs in a non-lethal muscular dystrophy context we generated *desma^+/-^ ;desmb^+/-^* zebrafish with premature stop codons in exon 1 of both genes (Fig. S1A-B). These mutations lead to truncated Desmin lacking α-helix-rod domains, essential for coil formation and tertiary structure formation. We confirmed that the *desma^-/-^;desmb^-/-^*double mutants, unlike *desma^+/-^;desmb^+/-^* controls, lacked Desmin immunolabeling at 3 dpf (Fig. S1C). Next, we performed functional experiments to investigate the impact of lack of Desmin in zebrafish. Larvae were reared in 1% methyl cellulose, to generate swimming resistance, between days 4 and 5. This triggered significant detachment and breaks, both in *desma^-/-^;desmb^-/-^* mutants and *desma^-/-^* single mutants, at the 10-12^th^ somite level (Fig. 1A-B), whereas *desmb^-/-^*and *desma^+/-^;desmb^+/-^* displayed no myofiber damage (Fig. 1B), indicating that *desma* is the main contributor to muscle tensile strength. Previous studies have shown both *desma* and *desmb* expression in developing zebrafish somite musculature^31^. Therefore, to investigate complete loss of *desmin* and avoid potential confounding factors, we decided to continue our studies using only *desma^-/-^;desmb^-/-^*zebrafish. Force/tension relationship measurements revealed that 5-6 dpf *desma^-/-^;desmb^-/-^* zebrafish larvae display a significant decrease in trunk myofiber maximal force generation when compared to *desma^+/-^;desmb^+/-^* (Fig. 1C-D). Furthermore, *desma^-/-^;desmb^-/-^* mutants showed a significant decrease in spontaneous movement in comparison to *desma^+/-^;desmb^+/-^*(Fig. 1E-F). This was not due to myofiber loss, as both genotypes had equal numbers of *Tg(mylz2:EGFP)* fast and *Tg(smyhc1:tdTomato)* slow twitch positive myofibers in cross sections of the trunk (Fig. S1D-E). Altogether, these results show that Desmin contributes to myofiber integrity, and is needed to maintain proper function of muscle tissue in the embryonic zebrafish trunk.

**Figure 1.**
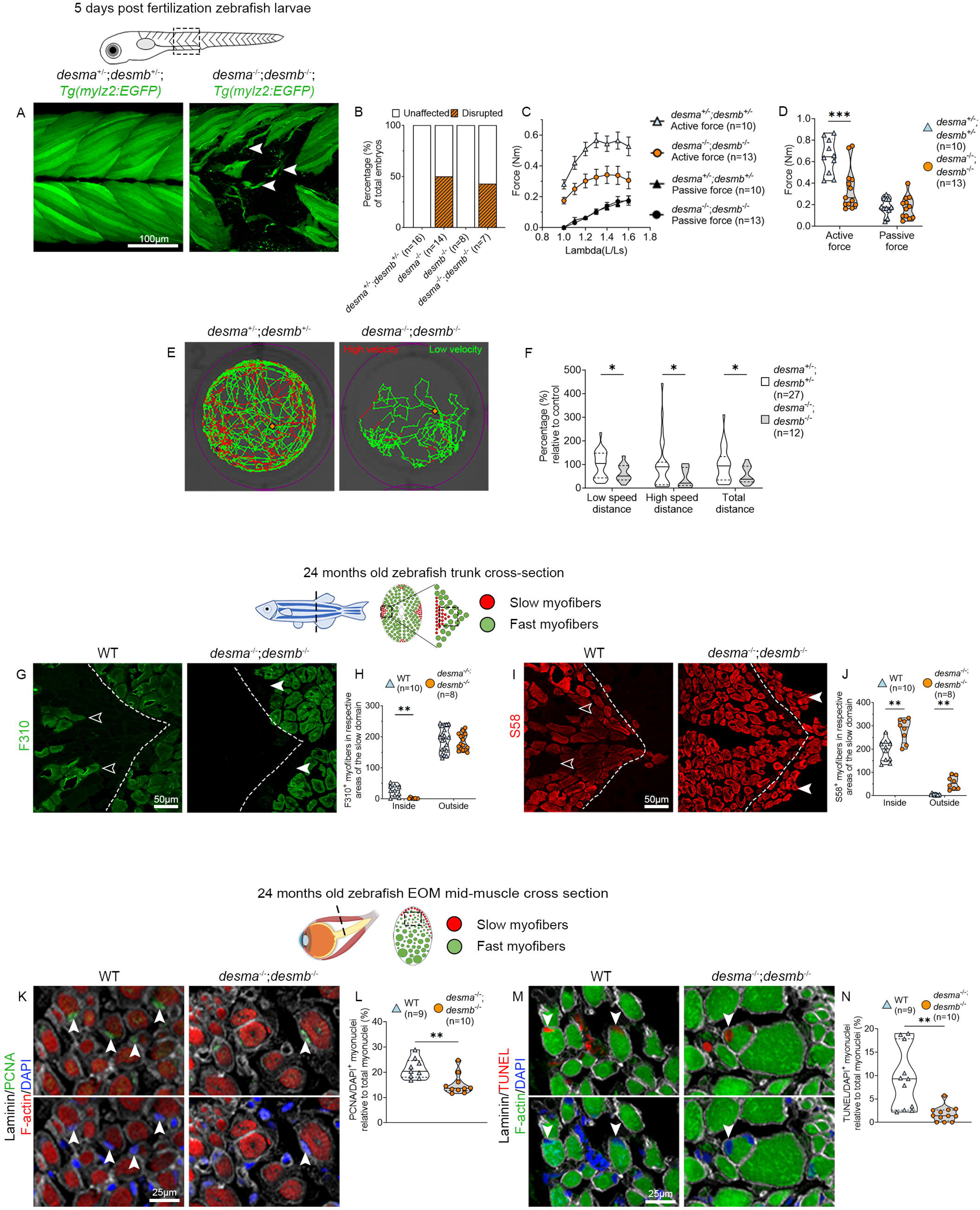
Lack of Desmin causes myofiber impairment and a glycolytic-oxidative metabolic shift in myofibers over time. A) Resistance swimming in 1% methylcellulose in embryo medium during overnight during 12 h from 4 to 5 dpf for *Tg(mylz2:EGFP);desma^-/-^ ;desmb^-/-^* mutant larvae and *desma^+/-^;desmb^+/-^*controls. B) Proportions of injuries caused by resistance swimming. C) Force generation of *desma^+/-^;desmb^+/-^*and *desma^-/-^;desmb^-/-^* trunk myofibers. D) Force generated at optimal stretch in *desma^+/-^;desmb^+/-^* and *desma^-/-^;desmb^-/-^*(p=0.002). E) Spontaneous swimming for 1 hour in *desma^-/-^;desmb^-/-^*and *desma^+/-^;desmb^+/-^* controls. F) Relative swimming activity at low speed (p=0.024), high speed (p=0.019) and total distance (p=0.016). G) Immunohistochemical analyses of tissue sections using F310 antibodies labeling fast myofibers in *desma^-/-^;desmb^-/-^* compared to WT (*desma^+/+^;desmb^+/+^*) controls. The separation of the fast and slow domains is defined by a myoseptum indicated by a dashed line. Open arrowheads indicate the presence of fast myofibers inside the slow muscle domain in WT. White arrowheads indicate lack of F310 labeling in the fast domain in *desma^-/-^;desmb^-/-^*. H) Comparison of the number of fast myofibers inside and outside of the slow domains respectively in WT versus *desma^-/-^;desmb^-/-^* (p=0.0018). I) Immunohistochemical analyses of tissue sections using S58 antibodies labelling slow myofibers in *desma^-/-^;desmb^-/-^* and WT controls. The separation of the fast and slow domains is defined by a myoseptum indicated by a dashed line. Open arrowheads indicate the lack of slow myofibers inside the slow muscle domain in WT. White arrowheads indicate presence of S58 labeling in the fast domain in *desma^-/-^;desmb^-/-^*. J) Comparison of the number of slow myofibers inside and outside of the slow domain respectively in WT versus *desma^-/-^;desmb^-/-^*. Inside (p=0.002), outside (p=0.0012). K) DAPI/PCNA/Laminin/Phalloidin immunolabeling of *desma^-/-^;desmb^-/-^* and WT EOMs. Arrowheads indicate DAPI/PCNA double positive myonuclei. L) Statistical analysis of DAPI/PCNA positive myonuclei inside the laminin sheet (p=0.0039). M) DAPI/TUNEL/Laminin/Phalloidin immunolabeling of *desma^-/-^;desmb^-/-^* and WT EOMs. Arrowheads indicate DAPI/TUNEL positive myonuclei. N) Statistical analysis of DAPI/TUNEL positive myonuclei inside the laminin sheet (p=0.0032). Data in violin plots (D, F, H, J, L, N) are presented as median (line) and quartiles (dashed lines). Data in C) are presented as mean ± SEM. Dashed lines in illustrations indicate the approximate level of the cross sections in figures.

### Extraocular muscle integrity is preserved in the zebrafish desminopathy model

Patients with desminopathy are often asymptomatic until mid-30s^29^. Therefore, to characterize our desminopathy model at later stages, we reared *desma^-/-^;desmb^-/-^*mutants and controls under equal conditions until 20-24 months of age, approximately two thirds of domesticated zebrafish life span, at which point they were histologically analyzed. *desma^-/-^ ;desmb^-/-^* mutants were fertile and survived until adulthood when reared separately from siblings (Fig. S1F). A consistent feature of muscular dystrophies is a glycolytic shift in myofiber identity from fast to slow^32,33^. To assess whether the lack of Desmin had an impact on myofiber identity, trunk and EOMs of 24 months old zebrafish were immunolabeled using antibodies against fast myosin light chain (F310) and against slow myosin heavy chain 1, 2 and 3 (S58)^34^. The zebrafish trunk consists of myofibers subdivided into compartments separated by thin layers of connective tissue, myosepta, easily identified on cross-sections (Fig. 1, dashed line in G, I). Slow myofibers are generally found in the most laterally positioned compartments, separated from fast myofibers by a hyperplastic growth zone positioned near the myosepta^35^. Adult zebrafish EOMs also display a distinct location of slow and fast myofibers, although not separated by a myoseptum^20^ (Fig. S1G, I). Cross-sections of trunk muscle of *desma^-/-^;desmb^-/-^* mutants revealed a decrease of fast F310 positive myofibers inside the slow domain (Fig. 1G-H, open arrowheads), a decrease in the slow myofiber diameter together with a significant increase in myofiber number in the slow domain, compared to WT controls (*desma^+/+^;desmb^+/+^*) (Fig. 1I-J). Additionally, the number of slow myofibers inside the fast-medial compartments was significantly increased (Fig. 1I-J, arrowheads). Cross-sections of EOMs showed that they remained unaffected in terms of proportion of fast and slow myofibers (Fig. S1G-J). In summary, adult *desma^-/-^;desmb^-/-^*mutant trunk muscle show a glycolytic-fast to oxidative-slow metabolic shift, consistent with a muscular dystrophy phenotype whereas the EOMs remain unaffected in this regard.

To further define the *desma^-/-^;desmb^-/-^* muscle phenotype, we analyzed zebrafish trunk muscle and EOMs for signs of regeneration, proliferation and cell death. *desma^-/-^;desmb^-/-^*mutant trunk muscle showed a significantly increased proportion of both Pax7 positive nuclei (Fig. S1K-L) and PCNA positive nuclei (Fig. S1M-N) in the slow myofiber domains compared to controls. No change in TUNEL positive nuclei in the slow domains was observed, however, the fast myofiber domains contained clusters of myofibers where most nuclei were TUNEL positive (Fig. S1O, arrows), along with a significant increase in centrally positioned nuclei, a hallmark of muscular dystrophy (Fig. S1O-P, asterisk). In contrast, controls only appeared to have sporadic signs of DNA fragmentation dispersed among nuclei in the entire myofiber population (Fig. S1O). Interestingly, in EOM cross sections of *desma^-/-^;desmb^-/-^* mutants, we instead noted a significant decrease of PCNA positive myonuclei (Fig. 1K-L) and TUNEL positive myonuclei (Fig. 1M-N) compared to controls, suggesting a decreased cellular turn over in response to the lack of Desmin in the EOMs. Overall, EOM myofiber composition remains unchanged despite the lack of Desmin, whereas trunk muscle is significantly affected and show clear signs of muscular dystrophy.

### Upregulation of fhl2b is an EOM response to muscular dystrophy

To investigate the adaptations observed in Desmin deficient EOMs we performed RNA-sequencing of EOMs and trunk muscle at 5 and 20 months of age, representing pre- and symptomatic stages, respectively. For each stage, we obtained transcriptome profiles from both trunk muscle and EOMs from *desma^-/-^;desmb^-/-^*mutant and WT controls (Fig. S2A, schematic illustration). To identify genes involved specifically in the EOMs resistance towards muscular dystrophy we performed differential expression analysis in *desma^-/-^;desmb^-/-^* compared to controls in EOMs (Fig. 2A, comparison I-II) and trunk (Fig. 2A, comparison III-IV) at both time points (Fig. S2A-D, Supplemental Table 1), additionally, we also compared EOMs to trunk (Fig. 2, comparisons V-VIII, Fig. S2E-H, Supplemental Table 1). Subsequently, we performed Gene Ontology (GO) analysis and retrieved differentially expressed genes (DEGs) contained within the cellular compartment GO terms related to myofiber function/structure (Fig. S3A-F, Supplemental Table 2), which we further cross-compared (Fig. 2A, comparisons I-VIII). Interestingly, when analyzing 20 months old EOMs we found upregulation of several DEGs related to cytoskeletal rearrangement (Fig. 2A, comparison III). One of these genes, *fhl2b,* was consistently identified across comparisons III-VIII highlighting it as a potential candidate gene (Fig. 2A). Notably, four members of the *fhl* family, including *fhl2b*, were identified in all EOM *vs* trunk comparisons (Fig. 2A). We did not find *fhl2b* to be differentially expressed in 5 months old *desma^-/-^;desmb^-/-^*EOMs which can likely be attributed to the early stage of disease progression. Next, we analyzed *fhl2b* gene expression across the different comparisons (Fig. 2A, comparisons I-VIII) and found that *fhl2b* consistently was more highly expressed in EOMs compared to trunk muscle (Fig. 2B). Previous studies have shown that *Fhl2* is mainly expressed in cardiac muscle^36^, localized to Z-discs, but has to our knowledge not been studied in the EOMs prior to this study. EOM longitudinal sections immunolabeled for Fhl2 showed localization to the Z-disc (Fig. S3G). The number of Fhl2-immunolabelled myofibers in EOMs of *desma*^-/-^;*desmb*^-/-^ mutants was significantly increased relative to controls at both 5 and 20 months of age and increased significantly with age (Fig. 2C-E) confirming our transcriptomic analysis. Next, we asked whether Fhl2 upregulation in EOMs could be found across muscular dystrophies and therefore immunolabeled *plecb^-/-^* (Plectin) and *obscnb^-/-^* (Obscurin) mutant zebrafish EOMs (Fig. 2F), both resulting in muscular dystrophy when mutated in other models^37,38^. Interestingly, *plecb^-/-^* and *obscnb^-/-^* mutant zebrafish displayed increased numbers of Fhl2 positive myofibers compared to controls, similar to *desma*^-/-^;*desmb*^-/-^ EOMs (Fig. 2C-F). This shows that Fhl2 is more widespread in several different cytoskeletal gene mutations and models for muscular dystrophy. Importantly, healthy adult human and mouse EOMs were also found to contain Fhl2 positive myofibers (Fig. 2G-H), indicating a putatively conserved role for Fhl2 in EOMs across species. In summary, Fhl2 is present in EOMs across different variants of muscular dystrophies and species and is increased with the progression of the disease in the EOMs of *desma*^-/-^;*desmb*^-/-^ mutant zebrafish.

**Figure 2.**
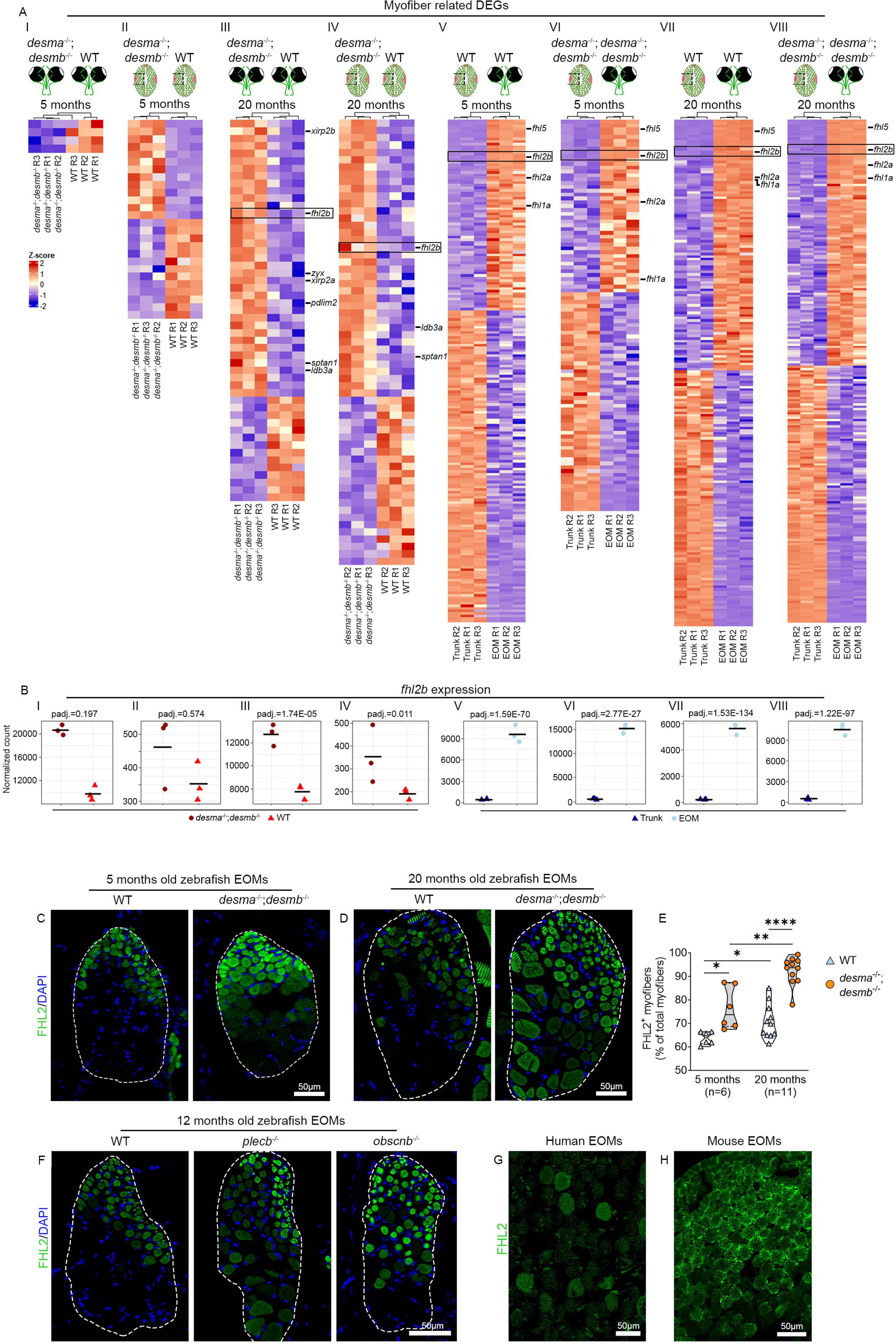
*fhl2b* is an EOM specific marker of EOM response to muscular dystrophy. A) Expression of myofiber-related DEGs for the following comparisons: 5 months *desma^-/-^ :desmb^-/-^ vs* WT EOMs (I), 5 months *desma^-/-^:desmb^-/-^ vs* WT trunk (II), 20 months *desma^-/-^ :desmb^-/-^ vs* WT EOMs (III), 20 months *desma^-/-^:desmb^-/-^ vs* WT trunk (IV), 5 months WT EOMs *vs* WT trunk (V), 5 months *desma^-/-^:desmb^-/-^* EOMs *vs desma^-/-^:desmb^-/-^* trunk (VI), 20 months WT EOMs *vs* WT trunk (VII) and 20 months *desma^-/-^:desmb^-/-^* EOMs *vs desma^-/-^ :desmb^-/-^* trunk (VIII). B) *fhl2b* expression in the abovementioned comparisons I-VIII. C) Fhl2 antibody labeling of WT and *desma^-/-^;desmb^-/-^* EOMs at 5 and D) 20 months. E) Fhl2 positive myofibers quantification between *desma^-/-^;desmb^-/-^* and WT control EOMs for 5 (p=0.017) and 20 months old zebrafish (p<0.0001), respectively. Quantification of 5 months versus 20 months old WT (p=0.014) and *desma^-/-^:desmb^-/-^* (p=0.005), respectively. Data is presented as median (line) and quartiles (dashed lines). F) WT, *plecb^-/-^*and o*bscnb^-/-^* 12 months old EOM cross section immunolabeling using Fhl2 antibodies. G) Human and H) mouse EOM cross section immunolabeling using Fhl2 antibodies. Scale bars: 50µm. White dashed lines outline the entire myofiber area of the EOMs.

### Knockout of fhl2 leads to EOM myofiber hypertrophy

Previous studies have found Fhl2 to be mainly expressed in cardiac muscle, however, *Fhl2* knockout mice were shown to be viable and no phenotype was observed unless challenged with isoproterenol, triggering adrenergic stimuli which resulted in cardiac hypertrophy^39^. We hypothesized that knockout of *fhl2* in the background of *desma^-/-^;desmb^-/-^*mutations could cause similar phenotypes in EOMs of adult zebrafish. We generated knockout lines of both zebrafish *fhl2* genes, *fhl2a* and *fhl2b*, to avoid possible confounding effects of redundancy (Fig. S4A-B). *In situ* hybridization of *fhl2a* showed low EOM expression whereas *fhl2b* was found to be distinctly expressed in the EOMs at 5 dpf (Fig. S4C-D, arrowheads) suggesting a larger role for *fhl2b* compared to *fhl2a* in the EOMs. Immunolabeling using Fhl2 antibodies confirmed lack of Fhl2 in *desma^-/-^;desmb^-/-^;fhl2a^-/-^;fhl2b^-/-^* compared to *desma^-/-^;desmb^-/-^ ;fhl2a^+/-^;fhl2b^+/-^* sibling controls (Fig. S4E).

Quantification of EOM myofiber size at 12 months of age in single *fhl2a^-/-^* and *fhl2b^-/-^* mutants and *fhl2a^-/-^;fhl2b^-/-^* double mutants, all in a *desma*^-/-^;*desmb*^-/-^ background, alongside the corresponding controls revealed that the myofiber areas were significantly increased in *desma^-/-^;desmb^-/-^;fhl2b^-/-^*and *desma^-/-^;desmb^-/-^;fhl2a^-/-^;fhl2b^-/-^*mutants compared to WT controls (Fig. 3A-B). This indicates that *fhl2b* has a role in hypertrophic protection, in line with results observed for *fhl2* in mouse cardiac muscle^39^. Given the reduction on cell death and proliferation observed in *desma^-/-^;desmb^-/-^*mutant EOMs (Fig. 1K-N), we wondered whether the knockout of *fhl2a* and *fhl2b* would revert these effects. For this purpose, we addressed cell death and proliferation on EOM cross sections. Myonuclei TUNEL (Fig. 3C-E) and PCNA (Fig. 3F-H) labeling showed a redundant relationship between *fhl2a* and *fhl2b*. Both *desma^-/-^ ;desmb^-/-^;fhl2a^-/-^* and *desma^-/-^;desmb^-/-^;fhl2b^-/-^* triple mutants displayed no change in cell death whereas a moderate increase in proliferation was found. However, in *desma^-/-^;desmb^-/-^;fhl2a^-/-^ ;fhl2b^-/-^* quadruple mutant zebrafish EOMs a highly significant increase in both cell death and proliferation was observed (Fig. 3E, H). These results indicate that *fhl2a* and *fhl2b* both maintain myonuclei integrity in the EOMs under muscle dystrophy conditions, however, *fhl2b* likely has an additional role in hypertrophic protection. Collectively, we show that *fhl2* is needed to maintain myofiber homeostasis in the EOMs in the background of *desmin* knockout and can therefore be considered as an EOM protective gene.

**Figure 3.**
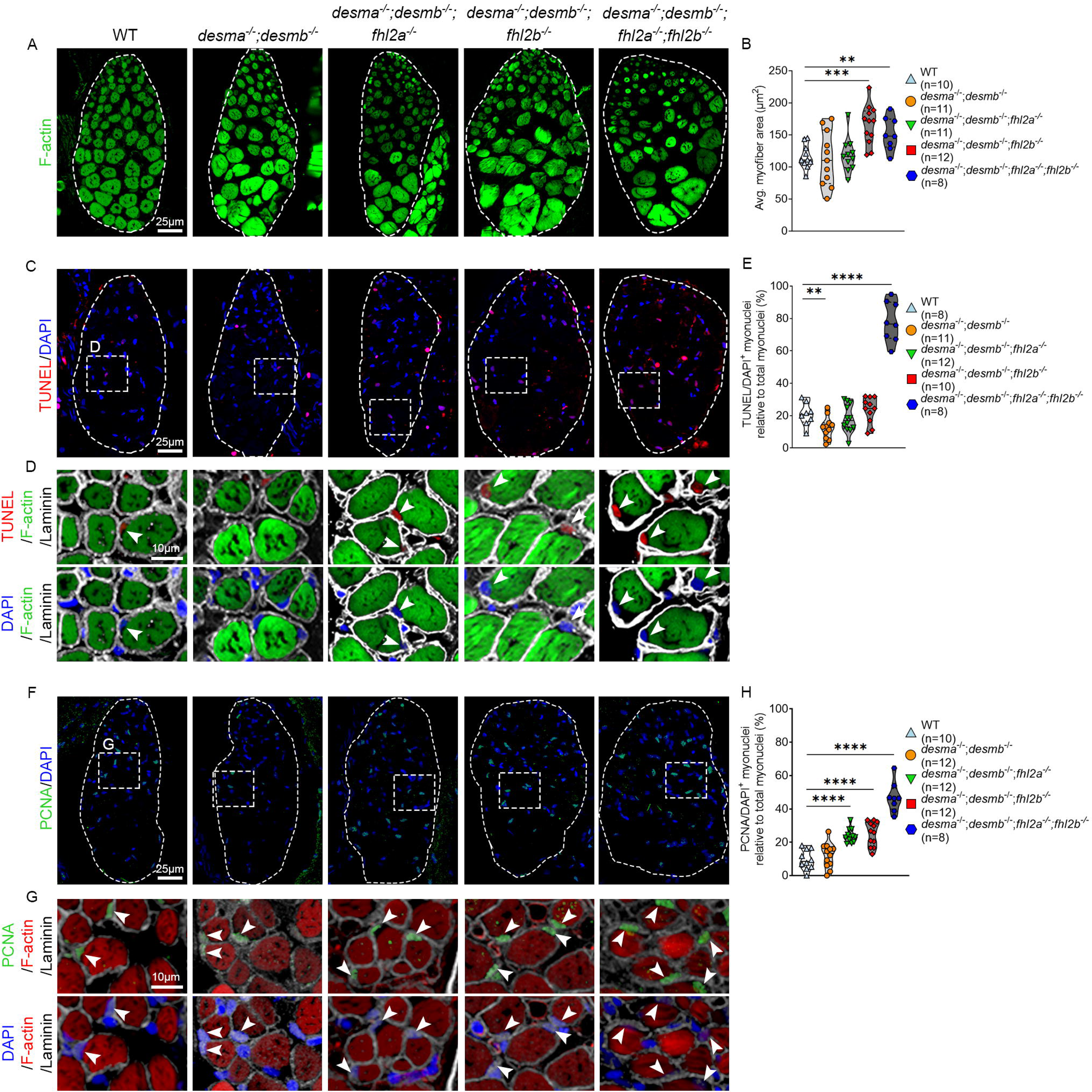
Lack of Fhl2 causes EOM myofiber hypertrophy. Cross-sections of 12 months old WT, *desma^-/-^;desmb^-/-^*, *desma^-/-^;desmb^-/-^;fhl2a^-/-^*, *desma^-/-^;desmb^-/-^;fhl2b^-/-^* and *desma^-/-^ ;desmb^-/-^;fhl2a^-/-^; fhl2b^-/-^* zebrafish EOMs. A) F-actin labeling (phalloidin) and B) myofiber area (F-actin positive) quantification showed significantly increased myofiber area in *desma^-/-^ ;desmb^-/-^;fhl2b^-/-^* (p=0.0002) and *desma^-/-^;desmb^-/-^;fhl2a^-/-^; fhl2b^-/-^* (p=0.007) mutant zebrafish. C) DAPI and TUNEL labeling of myonuclei. Dashed lines outline the entire myofiber area of the EOMs. Dashed boxes indicate magnified areas. D) Magnified areas indicated in C, DAPI and TUNEL labeling of myonuclei inside the laminin sheet. Phalloidin labels F-actin in the myofibers. White arrowheads indicate TUNEL positive myonuclei. E) DAPI/TUNEL positive myonuclei in *desma^-/-^;desmb^-/-^* were significantly reduced compared to WT (p=0.021) and significantly increased in *desma^-/-^;desmb^-/-^;fhl2a^-/-^; fhl2b^-/-^* compared to WT (p=<0.0001). F) DAPI/PCNA labeling of myonuclei. Dashed lines outline the entire myofiber area of the EOMs. Dashed boxes indicate magnified areas in G). G) DAPI/PCNA labeling of myonuclei inside the laminin sheet. Phalloidin labels F-actin in the myofibers. H) The number of PCNA positive myonuclei in *desma^-/-^;desmb^-/-^;fhl2a^-/-^*(p<0.0001), *desma^-/-^;desmb^-/-^;fhl2b^-/-^*(p<0.0001) and *desma^-/-^;desmb^-/-^;fhl2a^-/-^; fhl2b^-/-^* (p<0.0001) were significantly increased compared to WT. White arrowheads indicate DAPI/PCNA positive myonuclei. Data in all violin plots are presented as median (line) and quartiles (dashed line). Avg = average.

### Muscle specific overexpression of fhl2b significantly improves survival and myofiber integrity of the Duchenne muscular dystrophy zebrafish model

To investigate whether *fhl2b* also could protect muscles other than EOMs in muscular dystrophy and test this in a more severe context, we utilized the lethal zebrafish *sapje* (*dmd^ta222a^*) line^9^, a model of Duchenne muscular dystrophy, hereby referred to as *dmd^-/-^*. We overexpressed *fhl2b* under the muscle specific 503unc promoter^40^ coupled to EGFP via a T2A linker and analyzed trunk muscle in the *dmd^-/-^* background (Fig. 4A). We initially examined mosaic overexpression of *fhl2b*, which showed complete overlap between EGFP positive myofibers and Fhl2 antibody positive myofibers (Fig. S4F). *dmd^-/-^* larvae with a high number of EGFP positive myofibers generally showed less myofiber disruption than seen in *dmd^-/-^*larvae with low number EGFP positive fibers (Fig. S4G-H). We therefore reared mosaic larvae to adulthood and generated stable overexpression lines from three different founders with different EGFP intensity levels, one low and two high. To functionally test the effects of *fhl2b* overexpression on *dmd^-/-^*larvae, we analyzed survival among low and high level *fhl2b* expressing lines and found a significant correlation between *fhl2b* expression (Fig. S4I) and survival of *dmd^-/-^;Tg(503unc:fhl2b-T2A-EGFP)* larvae (Fig. 4B). The median survival time of *dmd^-/-^* larvae was found to be 18 dpf, whereas *dmd^-/-^;Tg(503unc:fhl2b-T2A-EGFP)* and sibling control median survival time was longer than 30 days (Fig. 4B). Notably, even low levels of *fhl2b* overexpression increased survival beyond 30 days in *dmd^-/-^;Tg(503unc:fhl2b-T2A-EGFP)*. Additionally, spontaneous swimming distance was also significantly increased in *dmd^-/-^;Tg(503unc:fhl2b-T2A-EGFP)* larvae compared to *dmd^-/-^* larvae at 5 dpf (Fig. 4C-D). Collectively, these results show that muscle specific *fhl2b* overexpression improves motor function and significantly prolongs the lifespan in this model of Duchenne muscular dystrophy.

**Figure 4.**
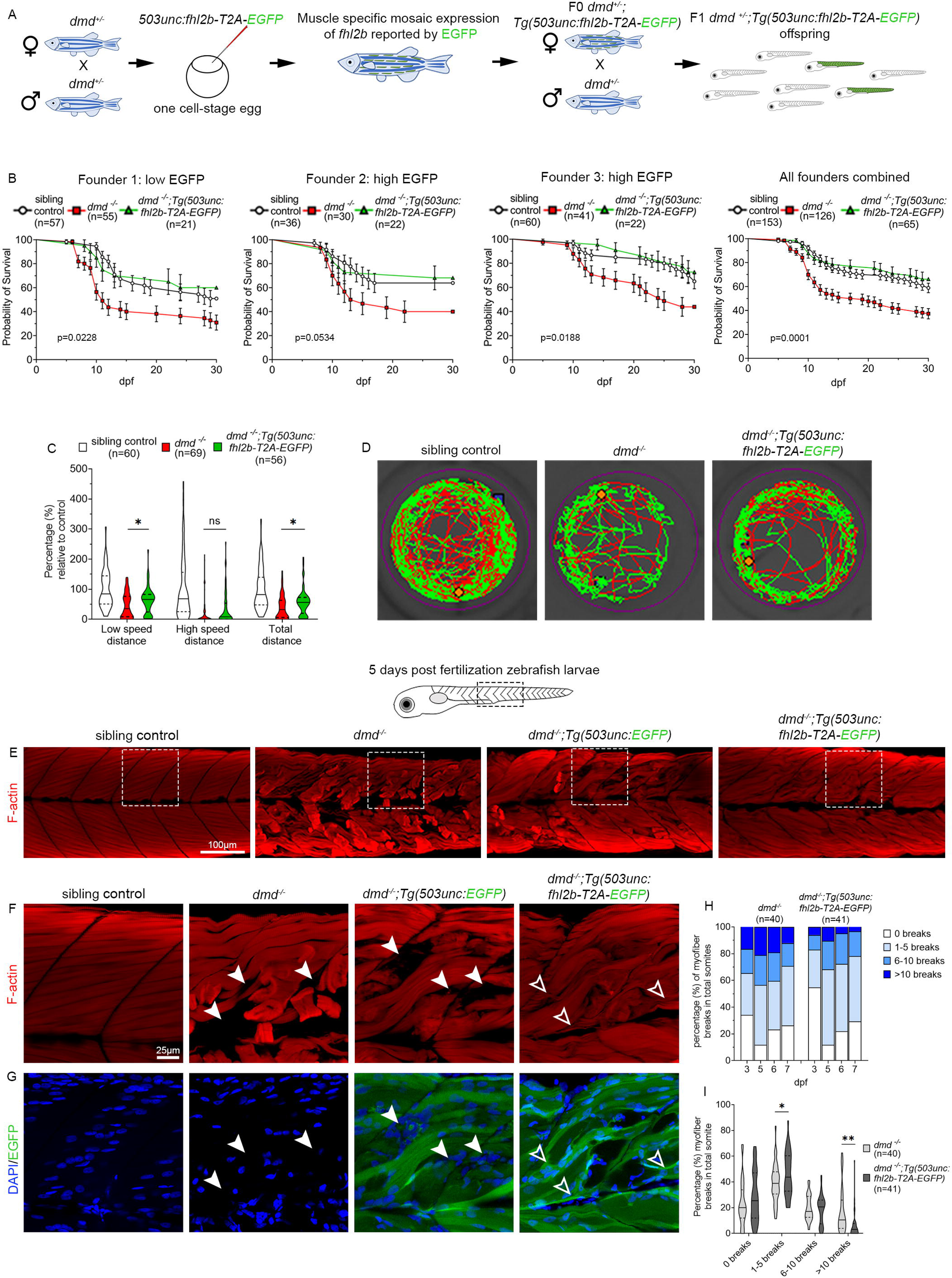
Muscle specific overexpression of *fhl2b* significantly prolongs life-span, improves motor function and muscle integrity in *dmd* zebrafish larvae. A) Experimental setup to generate *dmd^-/-^* zebrafish overexpressing *fhl2b*. One cell-stage eggs from in-crossed *dmd^+/-^* zebrafish were injected with a *503unc:fhl2b-T2A-EGFP* plasmid, raised and crossed into *dmd^+/-^* to generate stable lines. B) Survival tests over 30 days for three different *503unc:fhl2b-T2A-EGFP* lines. Kaplan-Meier log rank test was used to calculate significance between *dmd^-/-^*and *dmd^-/-^:Tg(503unc: fhl2b-T2A-EGFP)* for each of the three founder lines (Founder 1: p=0.0228, Founder 2: p=0.0534, Founder 3: p=0.0188 and all founder lines combined: p=0.0001). C) Spontaneous swimming tests performed over 1 hour showed significant increases in low speed (p=0.0459) and total distance (p=0.0385) in *dmd^-/-^ :Tg(503unc: fhl2b-T2A-EGFP)* compared to *dmd^-/-^* larvae. D) Representative swimming tracks of sibling control, *dmd^-/-^* and *dmd^-/-^:Tg(503unc: fhl2b-T2A-EGFP)* larvae. E) F-actin (phalloidin) labeling of trunk muscle in sibling control, *dmd^-/-^*, *dmd^-/-^:Tg(503unc:EGFP)* and *dmd^-/-^:Tg(503unc: fhl2b-T2A-EGFP)* larvae. F) Magnification of dashed boxes in (E) showing detached myofibers and empty areas in *dmd^-/-^*and *dmd^-/-^:Tg(503unc:EGFP)* larvae (arrowheads) whereas *dmd^-/-^:Tg(503unc: fhl2b-T2A-EGFP)* larvae display G) small diameter intensely EGFP positive myofibers (green) in corresponding areas (open arrowheads). DAPI in blue. H) Quantification of detached F-actin positive myofibers in somite segments at 3, 5, 6 and 7 dpf. I) Representation of number of myofiber breaks per somite. In total, *dmd^-/-^ :Tg(503unc: fhl2b-T2A-EGFP)* larvae show more small (1-5 breaks) myofiber detachments per somite (p=0.0441) and less large (>10 breaks) myofiber detachments per somite (p=0.001) as compared to *dmd^-/-^*larvae,. Data in violin plots (C, I) are presented as median (line) and quartiles (dashed line). Data in all survival graphs (B) are presented as mean ± SEM.

To examine if *fhl2b* overexpression improved myofiber integrity we labelled 5 dpf *dmd^-/-^*

*;Tg(503unc:fhl2b-T2A-EGFP)*, *dmd^-/-^* and sibling control (*dmd^+/+^*, *dmd^+/-^*) larvae using phalloidin and DAPI. Additionally, we generated a *Tg(503unc:EGFP)* line which was crossed into the *dmd^+/-^* line, used as a negative control. *dmd^-/-^;Tg(503unc:fhl2b-T2A-EGFP)* larvae showed detached myofibers similar to *dmd^-/-^* and *dmd^-/-^;Tg(503unc:EGFP)* larvae (Fig. 4E-F). However, *dmd^-/-^* and *dmd^-/-^;Tg(503unc:EGFP)* detached myofibers generated gaps devoid of F-actin positive myofibers in their direct proximity (Fig. 4F, closed arrowheads). Interestingly, detached myofibers in *dmd^-/-^;Tg(503unc:fhl2b-T2A-EGFP)* myotomes seldom had these gaps and thin F-actin positive myofibers were present (Fig. 4F, open arrowheads) which exhibited a high EGFP intensity, suggesting that they are newly formed as the unc503 promoter is more active in early stages of myogenesis^41^ (Fig. 4G). We quantified myofiber detachment in *dmd^-/-^* and *dmd^-/-^;Tg(503unc:fhl2b-T2A-EGFP)* larvae using F-actin. We found that *dmd^-/-^ ;Tg(503unc:fhl2b-T2A-EGFP)* consistently showed reduced number of myofiber detachments per somite compared to *dmd^-/-^* controls (Fig. 4H-I). Overall, these data indicate that *dmd^-/-^ ;Tg(503unc:fhl2b-T2A-EGFP)* myofibers have a lower tendency to detach compared to *dmd^-/-^* myofibers.

### Muscle specific fhl2b overexpression improves motor axon integrity and neuromuscular junctions in dmd^-/-^ larvae

To further understand how overexpression of *fhl2b* improves the *dmd^-/-^* phenotype, we analysed the transcriptomes of trunk muscle tissue from sibling controls (*dmd^+/+^*, *dmd^+/-^*), sibling*;Tg(503unc:fhl2b-T2A-EGFP), dmd^-/-^* and *dmd^-/-^;Tg(503unc:fhl2b-T2A-EGFP)* larvae using RNA-sequencing at 5 dpf (Fig. S5A-G, Supplemental Table 3). We then intersected the DEGs from the pairwise comparisons *dmd^-/-^ vs* sibling controls and *dmd^-/-^;Tg(503unc:fhl2b-T2A-EGFP) vs* sibling controls resulting in 1054 DEGs unique to *dmd^-/-^* larvae (Fig. 5A, gene set C). These 1054 DEGs correspond to *dmd* disease-related genes, whose expression is partially rescued by *fhl2b* overexpression (Fig. 5B). Gene ontology analysis of all intersections (Fig. S6A-B) revealed a unique enrichment for axon and neuron guidance-related terms in *dmd^-/-^*larvae (Fig. 5C), which notably included several *semaphorin* genes, crucial for axon guidance and motor neuron survival (Fig. 5D). To test whether axon and neuromuscular junction (NMJ) integrity were improved in *dmd^-/-^;Tg(503unc:fhl2b-T2A-EGFP)* larvae, we immunolabeled 5 dpf larvae for acetylated tubulin and α-bungarotoxin, labeling axons and post-synapse NMJs, respectively (Fig. 5E-G, Fig. S7). *dmd^-/-^* larvae displayed thin axons with reduced branching compared to sibling controls (Fig. 5E, G, arrowhead), additionally, NMJs were disorganized (Fig. 5F, G arrow) and an almost complete loss of contact was registered between axons and NMJs. In contrast, *dmd^-/-^;Tg(503unc:fhl2b-T2A-EGFP)* larvae displayed near normal axonal branching (Fig. 5E) and NMJ organization (Fig. 5G, open arrowhead). Our data indicate that axon retraction or degradation is a commonly occurring phenomenon in *dmd^-/-^* larvae. Furthermore, our data suggest that in myofibers rescued by *fhl2b*, axons and NMJs remain normal. However, we recognize that this may be the consequence of improved muscle tissue integrity.

**Figure 5.**
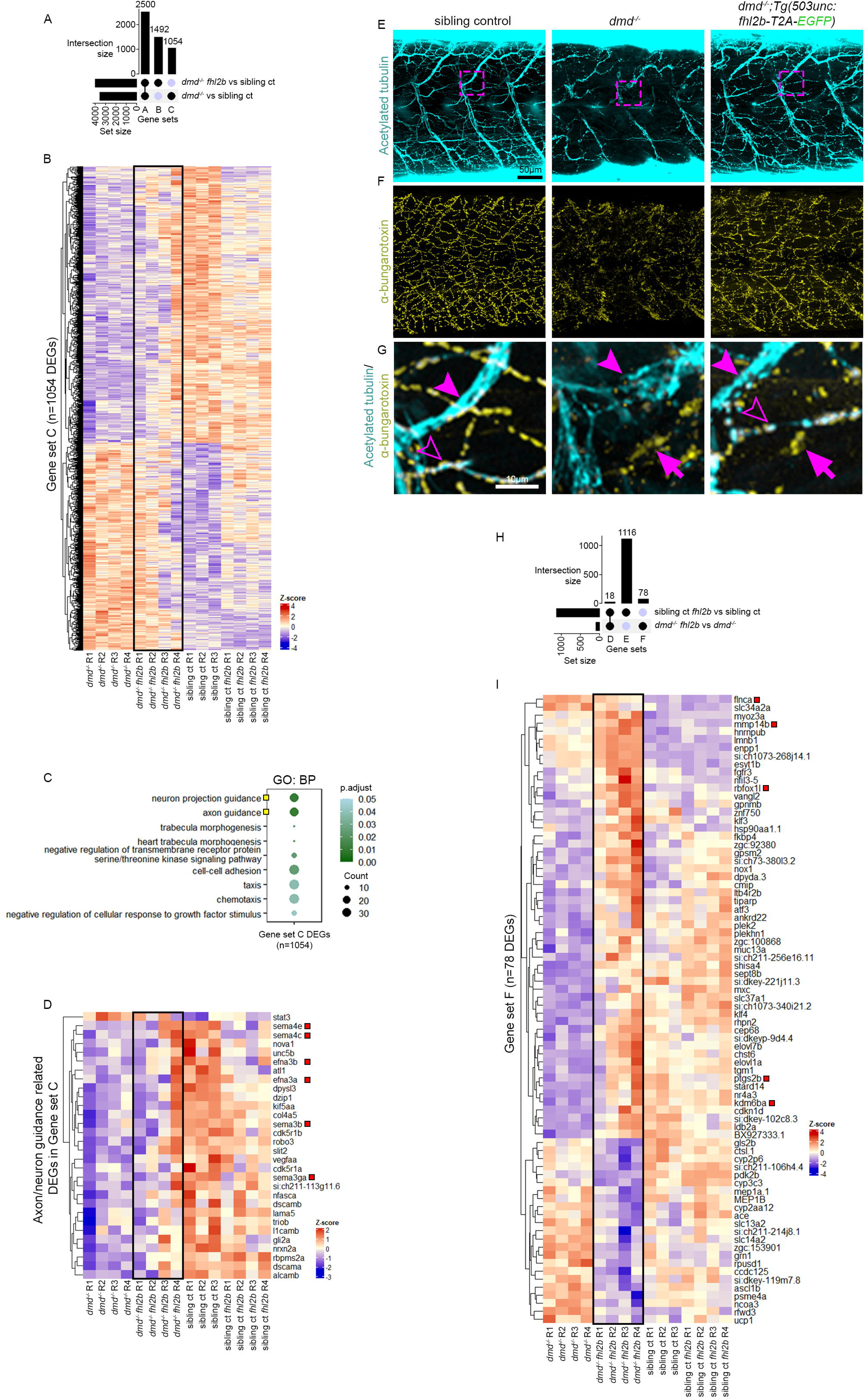
Muscle specific overexpression of *fhl2b* ameliorates axon and neuromuscular junction integrity in *dmd* zebrafish larvae. A) Upset plot showing the intersection of the DEGs between *dmd^-/-^:Tg(503unc: fhl2b-T2A-EGFP) vs* sibling controls (*dmd^+/+^*, *dmd^+/-^*) and *dmd^-/-^ vs* sibling controls (Gene set A: disease-related DEGs shared between *dmd^-/-^* and *dmd^-/-^:Tg(503unc: fhl2b-T2A-EGFP)* larvae; Gene set B: DEGs unique to *dmd^-/-^:Tg(503unc: fhl2b-T2A-EGFP)* larvae; Gene set C: disease-related DEGs specific to *dmd^-/-^* larvae). B) Heatmap displaying the expression of 1054 DEGs in Gene set C. C) Gene ontology terms enriched in genes from gene set C. Yellow squares highlight GO terms related to axon and neuron projection guidance; BP: Biological Process. D) Heatmap displaying the expression of axon and neuron guidance related DEGs, red boxes highlight *semaphorin* and *ephrin* DEGs. E) Sibling control, *dmd^-/-^* and *dmd^-/-^:Tg(503unc: fhl2b-T2A-EGFP)* larvae immunolabeled for Acetylated tubulin and F) α-bungarotoxin. G) Magnification of dashed boxes in E). Arrowheads indicate axons, open arrowheads indicate axon/NMJ overlap and arrows indicate NMJs lacking axon overlap. H) Upset plot showing the intersection of DEGs between *Tg(503unc: fhl2b-T2A-EGFP) vs* sibling controls and *dmd^-/-^ :Tg(503unc: fhl2b-T2A-EGFP) vs dmd^-/-^* larvae (Gene set D: DEGs caused by *fhl2b* overexpression in both comparisons; Gene set E: DEGs caused by *fhl2b* overexpression in healthy sibling control conditions alone; Gene set F: DEGs caused by *fhl2b* overexpression in the *dmd^-/-^* disease condition). I) Heatmap displaying the expression of DEGs in Gene set F. Red boxes indicate DEGs related to muscle regeneration.

### Muscle specific overexpression of fhl2b accelerates macrophage activity and muscle regeneration

To further define mechanisms underlying survival of *dmd^-/-^;Tg(503unc:fhl2b-T2A-EGFP)* larvae, we intersected DEGs from *dmd^-/-^;Tg(503unc:fhl2b-T2A-EGFP) vs dmd^-/-^* and sibling*; Tg(503unc:fhl2b-T2A-EGFP) vs* sibling controls (*dmd^+/+^*, *dmd^+/-^*) (Fig. 5H). We identified 78 genes differentially expressed in response to *fhl2b* overexpression in the *dmd^-/-^* disease background (Fig. 5H, gene set F). Among these 78 DEGs, we found several genes involved in muscle regeneration (Fig. 5I, *flnca*, *rbfoxl1, ptgs2b*, *kdm6ba*, *mmp14b*), suggesting that this is a possible mechanism for improved muscle integrity in *dmd^-/-^;Tg(503unc:fhl2b-T2A-EGFP)* larvae. In addition, we analysed 5 dpf *dmd^-/-^;Tg(503unc:fhl2b-T2A-EGFP)*, *dmd^-/-^*, *dmd^-/-^;Tg(503unc:EGFP)* and sibling controls for muscle regeneration, proliferation and cell death (Fig. S8A-G). Interestingly, we noted a significant decrease in Pax7 (Fig. S8A-B), BrdU (24 h pulse, Fig. S8C-E) and TUNEL (Fig. S8F-G) positive nuclei in *dmd^-/-^;Tg(503unc:fhl2b-T2A-EGFP)* compared to *dmd^-/-^* larvae, indicating overall healthier myofibers. These data confirm that *dmd^-/-^;Tg(503unc:fhl2b-T2A-EGFP)* myofibers are partially rescued by *fhl2b* overexpression and support our findings that *dmd^-/-^;Tg(503unc:fhl2b-T2A-EGFP)* myofibers are less prone to detachment. Interestingly, our study indicates that *fhl2b* induces faster formation of new myofibers and we therefore hypothesized that the wound healing process would be quicker in *fhl2b* overexpressing larvae. To address this, we performed timelapse studies on *dmd^-/-^;Tg(503unc:EGFP)* and *dmd^-/-^;Tg(503unc:fhl2b-T2A-EGFP)* larvae during 24 h to observe muscle regeneration in real time. We observed that detached myofiber debris was quickly removed and myofibers were apparently replaced in *dmd^-/-^;Tg(503unc:fhl2b-T2A-EGFP)* larvae whereas practically no change was observed in *dmd^-/-^;Tg(503unc:EGFP)* controls during the 24 h window (Fig. 6A). To quantify this, we measured birefringence in wounded somites from 3-5 dpf in *dmd^-/-^;Tg(503unc:fhl2b-T2A-EGFP)* and *dmd^-/-^ ;Tg(503unc:EGFP)* controls and found significantly improved birefringence in wounded somites in *dmd^-/-^;Tg(503unc:fhl2b-T2A-EGFP)* over time as compared to *dmd^-/-^ ;Tg(503unc:EGFP)* controls (Fig. 6B-C). Collectively, these results suggest that *fhl2b* overexpression accelerates muscle regeneration.

**Figure 6.**
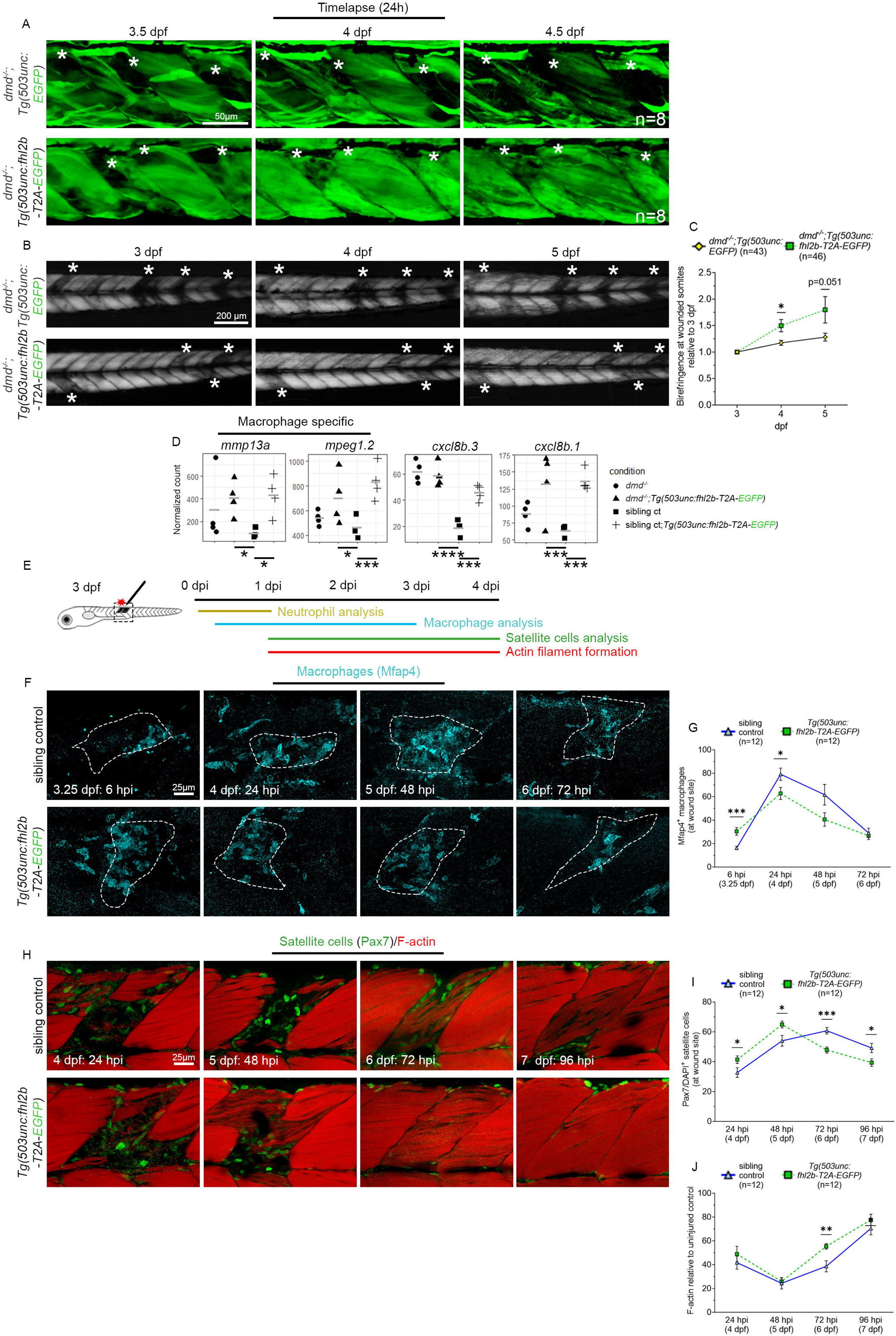
Muscle wound healing is enhanced in *fhl2b* overexpressing larvae. A) Timelapse images of *dmd^-/-^;Tg(503unc:EGFP)* and *dmd^-/-^;Tg(503unc:fhl2b-T2A-EGFP)* larvae at 3.5, 4 and 4.5 dpf respectively. Asterisks indicate damaged somites. B) Birefringence images following the same *dmd^-/-^;Tg(503unc:EGFP)* and *dmd^-/-^ ;Tg(503unc:fhl2b-T2A-EGFP* larvae at 3, 4 and 5 dpf. Asterisks indicate damaged somites. C) Quantification of birefringence in damaged somites over time. Birefringence at 4 and 5 dpf was compared to 3 dpf in order to evaluate wound healing progression over time. D) Normalized counts for *mmp13a* (p.adj=0.012, padj=0.023), *mpeg1.2* (p.adj=0.015, padj=0.0009), *cxcl8b.3* (p.adj=7.4e^-8^, padj=0.0007) and *cxcl8b.1* (p.adj=0.0005, padj=0.0001). Comparisons were made between *dmd^-/-^:Tg(unc503: fhl2b-T2A-EGFP) vs* sibling controls (*dmd^+/+^*, *dmd^+/-^*) and sibling*:Tg(unc503: fhl2b-T2A-EGFP) vs* sibling controls, respectively. E) Wound healing assay experimental setup. F) Macrophage specific Mfap4 immunolabeling of sibling controls and *Tg(unc503: fhl2b-T2A-EGFP)* between 6-72 hpi. Dashed lines indicate wounded areas. G) Quantification of Mfap4/DAPI positive cells at 6 (p=0.0019), 24 (p=0.0374), 48 and 72 hpi. H) Satellite cell specific Pax7 immunolabeling and F-actin of sibling controls and *Tg(unc503: fhl2b-T2A-EGFP)* at 24-96 hpi. I) Quantification of Pax7/DAPI positive cells in sibling controls and *Tg(unc503: fhl2b-T2A-EGFP)* at 24 (p=0.0445), 48 (p=0.0198), 72 (p=0.0002) and 96 hpi (p=0.0246). J) Quantification of F-actin intensity at wound site in sibling controls and *Tg(unc503: fhl2b-T2A-EGFP)* at 24, 48, 72 (p=0.0045) and 96 hpi. Data in graphs are presented as mean ± SEM.

Given our results showing that detached myofibers quickly disappeared in *dmd^-/-^ ;Tg(503unc:fhl2b-T2A-EGFP)* larvae (Fig. 6A), we hypothesized that increased leucocyte activity could be involved in the wound healing process. Leucocytes have previously been shown to be critical in zebrafish muscle regeneration^42^. Additionally, macrophage specific genes *mmp13a* and *mpeg1.2* as well as *il-8* were upregulated in *dmd^-/-^;Tg(503unc:fhl2b-T2A-EGFP)* and uninjured sibling*;Tg(503unc:fhl2b-T2A-EGFP)* as compared to sibling controls (Fig. 6D), supporting this hypothesis. To address this, we analyzed *fhl2b*’s effect on various aspects of muscle regeneration in a controlled setting using *Tg(503unc:fhl2b-T2A-EGFP)* larvae compared to sibling controls. We performed needle-stick injuries and analyzed the muscle regeneration process by immunolabeling for neutrophils (Mpx), macrophages (Mfap4) and satellite cells (Pax7) coupled with DAPI and phalloidin. All three antibodies were found to label an equal number of cells in 3 dpf uninjured controls (Fig. S.8H-M), and the neutrophil population labelled by Mpx antibodies was unchanged across all stages tested, from uninjured to 24 hours post injury (hpi) (Fig. S8N-O). Noticeably, we found a significant increase in macrophage count at the wound site in *Tg(503unc:fhl2b-T2A-EGFP)* larvae already at 6 hpi (Fig. 6E-G). This was followed by a significantly reduced number of macrophages at 24 hpi in *Tg(503unc:fhl2b-T2A-EGFP)* wounds compared to sibling control wounds (Fig. 6E-G). Pax7 positive satellite cells were present at wound sites faster in *Tg(503unc:fhl2b-T2A-EGFP)* compared to sibling controls and peaked at 48 hpi whereas sibling control satellite cell numbers peaked at 72 hpi (Fig. 6H-I). Additionally, F-actin intensity was found to be significantly higher at 72 hpi as compared to controls (Fig. 6J). Taken together, these results suggest that *Tg(503unc:fhl2b-T2A-EGFP)* larvae have faster myofiber regeneration via enhanced macrophage recruitment.

## Discussion

In the current study we show that induced expression of *fhl2b* can provide resistance to muscular dystrophy and that the EOMs offer a novel approach regarding protective cellular strategies. Here we propose a model (Fig. 7) in which *dmd^-/-^* larvae exhibit prolonged survival due to protection by *fhl2b*, muscle injury does not occur as frequently or to the same extent and axons and neuromuscular junctions are less affected. Additionally, *fhl2b* improves macrophage response to myofiber injury, thereby accelerating the process. We therefore propose that *fhl2b* based therapy can be utilized to alleviate muscular dystrophy in body musculature.

**Figure 7.**
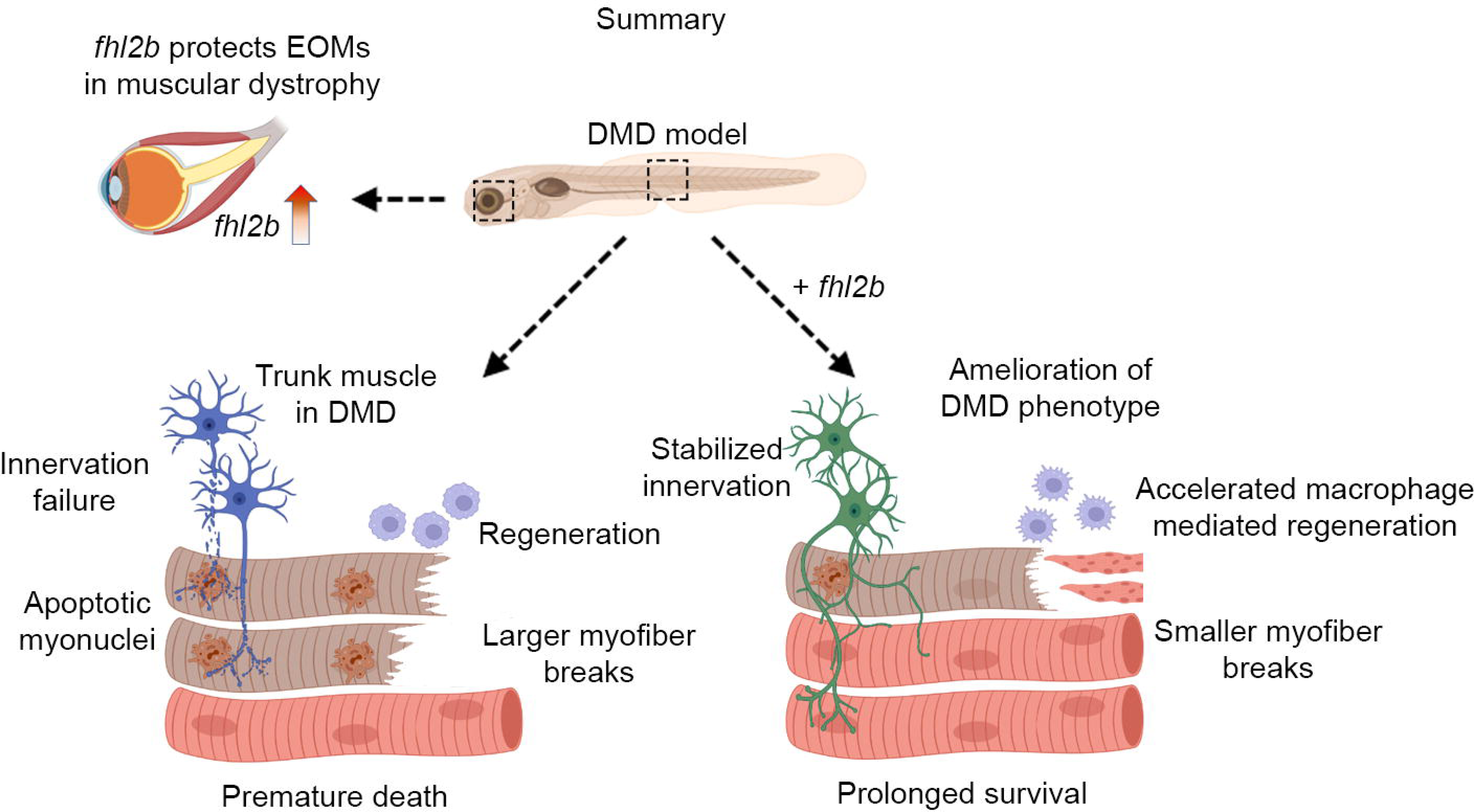
Model for *fhl2b* mediated rescue of muscular dystrophy. *Fhl2b* is upregulated and protects EOMs from muscular dystrophy conditions. *dmd^-/-^* larvae overexpressing *fhl2b* survive for extended amounts of time due to improved muscle integrity with fewer myofiber detachments and breaks, improved axon and NMJ stability and accelerated myofiber regeneration. Image generated by BioRender.

Fhl2 is a co-transcription factor and can translocate to the nucleus to co-activate gene expression^43^. However, we did not detect any Fhl2 positive myonuclei in the EOMs, instead Fhl2 was present mainly on the Z-discs of myofibers and likely executes its function there. In cardiomyocytes, Fhl2 has been shown to block ERK2 translocation to the nucleus and subsequent transcriptional activity, offering protection from hypertrophy^44^. Furthermore, Fhl2 has been linked to metabolic enzyme targeting to the N2B region of titin, suggesting that it acts as an adaptor protein for high energy demanding regions within the sarcomere^45^. The N2B region has also been proposed to be an anchoring hub for mechanotransduction^46^ and Fhl2 has previously been shown to be important for mechanotransduction in several scenarios where it is situated on F-actin and can translocate to the nucleus upon a change of tension to regulate gene expression^43,47^. Interestingly, *Zyx, pdlim2* and *ldb3a*, identified in our transcriptional profiling of dystrophic EOMs, are also linked to mechanotransduction^48-50^, introducing mechanosensing as a potentially important mechanism for EOM specific resistance to muscular dystrophy. This notion is strengthened by the fact that we found Fhl2 to be more abundant in a number of muscular dystrophy models, suggesting it is part of a general mechanism of defense in the EOMs. Knockout of *fhl2* leads to increased levels of cellular turn around and hypertrophy, emphasizing the importance of *fhl2* in EOM myofiber homeostasis, similar to what has been shown in rodent cardiomyocytes^39,44^. We show that ectopic *fhl2b* expression in *dmd^-/-^* larvae leads to protection from myofiber integrity loss. However, whether ectopic *fhl2b* has the same mechanism in trunk muscle, EOMs and cardiomyocytes remains to be determined.

Duchenne muscular dystrophy models have shown alterations in NMJ patterning^51^, non-surprising considering the dystrophin-glycoprotein-complex (DGC) normal accumulation at focal adhesion sites. Interestingly, our RNA-sequencing and *in vivo* data clearly demonstrate preserved expression of several genes related to axon and motor neuron guidance and a clear improvement in axon and NMJ structure in *dmd^-/-^;Tg(503unc:fhl2b-T2A-EGFP)* compared to *dmd^-/-^* larvae. *fhl2b* overexpressing larvae notably showed higher levels of axon guidance cues such as semaphorins, one of the major families of axon guidance molecules^52^. While muscle secreted molecules have been shown to affect motor neurons directly^53^, it is difficult to assess whether our results are a direct effect of *fhl2b* overexpression or a consequence of preserved myofiber integrity. Denervated myofibers undergo degeneration^54^, likely adding to the overall muscle phenotype observed in *dmd^-/-^* larvae. Previous studies have shown beneficial effects of electrical muscle stimulation on DMD deficient myofibers^55^. In our study, *fhl2b* overexpressing *dmd^-/-^* larvae showed improved motor function, which is likely a consequence of both improved muscle and axon/NMJ structure.

Fhl2 is linked to wound healing in several tissues^56,57^ and lack of Fhl2 or its suppression in these processes is coupled with chronical inflammation or deteriorating wound healing^58,59^. In muscle, Fhl2 has been suggested to regulate *il-6* and *il-8* production via MAPK signaling, playing a role in post-injury inflammation^60,61^. Our data show increased *il-8* levels under uninjured and *dmd^-/-^ fhl2b* overexpression conditions, supporting these claims. Recently, macrophages have been shown to be critical in muscle injury regeneration by inducing satellite cell proliferation^42^. Interestingly, our RNA-sequencing data show upregulation of macrophage specific genes (*mmp13a*, *mpeg1.2*) in uninjured *fhl2b* overexpressing sibling and *dmd^-/-^* larvae. Furthermore, in needle-stick injured *fhl2b* overexpressing sibling larvae, macrophages are significantly more abundant in the early stage of myofiber repair and regeneration compared to controls. Subsequently, the number of Pax7 positive satellite cells is significantly higher at an early stage of regeneration. As a result, myofibers are restored more rapidly at the wound site. In summary, we propose that overexpression of *fhl2b* results in accelerated muscle regeneration via enhanced macrophage recruitment.

Taking advantage of the EOMs innate resistance to muscular dystrophies combined with genetic models constitutes a novel approach to identify treatment strategies for these devastating conditions. Our study shows Fhl2 upregulation in EOMs of several disease models, suggesting that Fhl2 is a strong candidate for the development of future therapeutic strategies for a common management of muscular dystrophies.

## Author contributions

ND, JvH and FPD designed the study. Experiments were performed, analyzed and interpreted by ND, AK, IN, MC, HN, JL, AA and LJB. Transcriptional data was analyzed and interpreted by ND and IN, with supervision from SR. ND wrote the manuscript with supervision and input from LJB, SR, JvH and FPD. FPD secured funding for the project.

Corresponding author: Fatima Pedrosa Domellöf, Jonas von Hofsten.

## Acknowledgements

The authors acknowledge support from the National Genomics Infrastructure in Stockholm and Uppsala funded by Science for Life Laboratory, the Knut and Alice Wallenberg Foundation and the Swedish Research Council, and SNIC/Uppsala Multidisciplinary Center for Advanced Computational Science for assistance with massively parallel sequencing and access to the UPPMAX computational infrastructure. The computations were enabled by resources in projects SNIC 2022/22-442 and NAISS 2023/22-287 provided by the Swedish National Infrastructure for Computing (SNIC) and the National Academic Infrastructure for Supercomputing in Sweden (NAISS) at UPPMAX. The authors acknowledge Philip W. Ingham for generously sharing the *Tg(smyhc1:tdTomato)^i261^*and *Tg(mylz2:EGFP)^i135^* zebrafish lines. AA was funded by grants from Hans-Gabriel and Alice Trolle-Wachtmeister’s Foundation for Medical Research. JvH was funded by biotechnology grant for basic science FS 2.1.6-1911-22, the Medical Faculty, Umeå University. FPD was supported by research grants from the Swedish Research Council (Dnr 2018-02401), the Västerbotten County Council (Central ALF and Spjutspetsmedel), Kronprinsessan Margaretas Arbetsnämnd för synskadade (Stiftelsen KMA), Ögonfonden. ND was supported by Kempestiftelserna, Ögonfonden and Arnerska forskningsfonden.

## Experimental procedures

### Animal husbandry and ethical approval

Zebrafish larvae (*Danio rerio*) and adult fish were maintained from AB WT. Mutant lines used were *desma*^umu10^, *desmb*^umu11^, *obscnb^umu16^*, *plecb^umu25^*, *fhl2a^umu32^*, *fhl2b^umu33^*and *sapje^t222a^* (referred to as *dmd*^+/-^). Transgenic lines used were *Tg(mylz2:EGFP)^i135^*, *Tg(smyhc1:tdTomato)^i261^*, *Tg(503unc:EGFP)^umu37^* and *Tg(503unc:fhl2b-T2A-EGFP)^umu34^*. All zebrafish of the same genotype were reared from the same parental couple, to minimize genetic background bias across age. Additionally, WT zebrafish used originated from the same line utilized when generating the *desma^-/-^;desmb^-/-^* mutant. Zebrafish were maintained by standard procedures on a 10/14h dark/light cycle at 28°C at the Umeå University Zebrafish Facility. All animal experiments were ethically approved by the Regional Ethics Committee at the court of Appeal of Northern Norrlands Umeå djurförsöksetiska nämnd, Dnr: A6 2020.

### Generation of *desma*, *desmb*, *fhl2a* and *fhl2b* mutant zebrafish using CRISPR/Cas9

*desma* and *desmb* zebrafish mutants were generated using methods previously described^62^. For *desma*, and *desmb* a guide RNA (gRNA) targeting exon 1 was synthesized using the sequences 5’-ATTCAGCCTCCGCCGAGTCGG-3’ and 5’-GGTGGGTCGGGCAGCTCTCGG-3’ respectively, and was coupled with a gRNA scaffold^62^. The gRNA was then transcribed using the MegaShortScript T7 (Invitrogen) kit and co-injected with Cas9 protein (New England Biolabs) into one-cell stage zebrafish eggs. Injected larvae were grown to adulthood, outcrossed into WT zebrafish and screened to identify founders containing germline mutations. Mutant zebrafish larvae carrying a 5 bp and a 20 bp deletion in the *desma* and *desmb* genes, respectively, were chosen for further examination. Genotyping of *desma* mutant larvae was accomplished by standard PCR protocols using forward 5’-ATAGAAGTGGGCGCCAATG-3’ and reverse 5’-GTCTTGAGGAGCCAGAGGAA-3’ primers and *desmb* mutant larvae were genotyped using forward 5’-AGCCACTCTTATGCCACCTC-3’ and reverse 5’GCGGTCATTTAGATGCTGAAG-3’ primers. The PCR products were then digested overnight using HinfI and AluI for *desma* and *desmb*, respectively, and analyzed on a 2% agarose gel. *fhl2a* and *fhl2b* mutants were generated as described above, using gRNA sequences 5’-AAGAAGTATGTCCTGCGTGAGG-3’ and 5’-CGGGAAGAAGTACGTCCTGCGG-3’ respectively. To genotype *fhl2a* mutants forward 5’ATAGAAGTGGGCGCCAATG-3’ and reverse 5’TGGGTTTCTTGCATTCCTCG-3’ primers were used and the resulting PCR product was digested with ecoNI to screen for mutations. *Fhl2b* mutants were genotyped using 5’CCCTTTCAACCTGTCTGCAC-3’ and reverse 5’GCAGGTATTGGAGTAGAGGCT-3’ primers and the PCR product was digested using HpyCH4IV. The resulting *fhl2a* and *fhl2b* mutants chosen for investigation both carried 7 bp deletions in exon 1.

### Transgenesis

An overexpression vector (*503unc:fhl2b-T2A-EGFP*) containing the muscle specific promoter 503unc driving expression of *fhl2b* coupled to EGFP via T2A was acquired from Vectorbuilder (Neu-Isenburg, Germany). Subsequently, AgeI and NheI was used to remove the *fhl2b* cassette and fused using ligase, to generate a *503unc:EGFP* vector. Both vectors were co-injected in one cell-stage AB WT zebrafish eggs with Tol2-transposase RNA, at 30 ng/ul, respectively. Founders were screened using EGFP expression using fluorescent dissecting microscopy and out crossed with wild type fish to generate stable lines used in our experiments.

### Immunohistochemistry and TUNEL assay

Adult zebrafish at the appropriate age were deeply anesthetized with 0.01% ethyl 3-aminobenzoate methane sulfonate (Tricaine, MS-222, Sigma Aldrich) and sacrificed by decapitation. The heads and posterior trunks were fixed in 2% paraformaldehyde (PFA) for 1 hour at RT and stepwise incubated in 10, 20 and 30% sucrose for 12 hours each, at 4°C. The heads and trunks were then mounted separately on cardboard using OCT cryomount and snap frozen in liquid nitrogen chilled propane (-160°C), and finally cut serially into 12µm (trunk tissue) or 14µm (head tissue) thick sections using a cryostat (Reichert Jung; Leica, Heidelberg, Germany). Trunk preparations were cut from adult zebrafish at the anal opening and cut again 8-10mm caudally to the initial cut. Sections were always made from the proximal end of the specimen to assure the highest possible section similarity between fish. Sectioned zebrafish muscle tissue was rinsed in phosphate buffered saline (PBS) for 15 min before the addition of blocking solution (1% blocking reagent (Roche Diagnostics GmBH, Manheim, Germany) with 0.4% Triton X, 5% dimethyl sulfoxide (DMSO) and 0,1 % TWEEN20) for 1 hour, at RT. Primary antibodies were diluted in blocking solution and were applied for 48 hours, at 4°C. Sections were then washed 3 times in PBS and incubated with secondary antibodies for 24 h at 4°C, washed 3 times in PBS and mounted using 80% glycerol.

For whole mount immunohistochemistry, larvae were fixed in 2% PFA for 1 hour at RT, washed 3 times in PBS, acetone cracked for 1 hour at -20°C, washed 3 times in PBS and incubated for 1 hour with blocking solution at RT. Blocking solution was replaced by blocking solution containing primary antibody and incubated from 48-96 hours depending on stage. Larvae were washed 3 times 15 min in PBS before addition of secondary antibodies diluted in blocking solution and incubated 24-48 hours depending on stage before being washed 3 times 15 min in PBS again. Lastly, larvae were allowed to equilibrate to 80% glycerol for 1 hour before mounted on glass slides for imaging. DAPI and phalloidin was added with the secondary antibodies. All primary and secondary antibodies used are presented in Table 1.

**Table 1.** Antibody list.

| **Primary antibody** | **Host** | **Dilution** | **Source** | **Productnr/clone** |
| --- | --- | --- | --- | --- |
| F310 | Mouse | 1;10 | DSHB | AB_531863 |
| S58 | Mouse | 1;10 | DSHB | AB_528377 |
| Pax7 | Mouse | 1;10 | DSHB | AB_528428 |
| Desmin | Rabbit | 1;100 | Abcam | ab15200 |
| Laminin | Rabbit | 1;100 | Sigma | L9393 |
| PCNA | Rabbit | 1;100 | Sigma | SAB2701819 |
| Acetylated Tubulin | Rabbit | 1;100 | Sigma | T7451 |
| FHL2 | Rabbit | 1;100 | Sigma | HPA005922 |
| MPX | Rabbit | 1;100 | GeneTex | GTX128379 |
| MFAP4 | Rabbit | 1;100 | GeneTex | GTX132692 |
| **Secondary antibodies** | **Against** | **Dilution** | **Source** |  |
| Alexa Fluor 488 | donkey anti rabbit IgG | 1;300 | Invitrogen, Molecular probes | A-21206 |
| Alexa Fluor 488 | donkey anti-mouse IgG | 1;300 | Invitrogen, Molecular probes | A-21202 |
| Rhodamine Red™-X | donkey anti rabbit | 1;500 | Jackson ImmunoResearch | AB_2340613 |
| Rhodamine Red™-X | donkey anti-mouse | 1;500 | Jackson ImmunoResearch | AB_2340832 |
| Alexa Fluor 647 | goat anti-mouse IgG | 1;300 | Invitrogen, Molecular probes | A32795 |
| Alexa Fluor 647 | goat anti-rabbit IgG | 1;300 | Invitrogen, Molecular probes | A32787 |
| Alexa Fluor 488 Phalloidin | Directly conjugated | 1;100 | Invitrogen, Molecular probes | A12379 |
| Phalloidin rhodamine | Directly conjugated | 1;100 | Invitrogen, Molecular probes | 899165 |
| Alexa Fluor 647 Phalloidin | Directly conjugated | 1;100 | Invitrogen, Molecular probes | A22287 |
| DAPI | Directly conjugated | 1;500 |  |  |
| BrdU-555 | Directly conjugated | 1;500 | BD Biosciences | 560210/clone 3D4 |

Additionally, a TUNEL assay (Click-iT, Alexa Flour, Invitrogen, Thermo Fisher Scientific) was performed to label nuclei containing degraded DNA molecules, following the manufacturer’s description.

### Whole-mount *in situ* hybridization

RNA probes for *in situ* hybridization were synthesized using a PCR method for RNA probes as previously described^63^. Zebrafish larvae were fixed in 4% paraformaldehyde (PFA) overnight at 4 °C and dehydrated in 30%, 50%, 70% and 100% methanol and stored in 100% methanol at -20 °C until use. Whole-mount *in situ* hybridization was performed as described previously^64^. The RNA probes were based on the sequences of *fhl2a* (NM_001003732.1) and *fhl2b* (NM-001006028.2). Primers used to generate probes were: *fhl2a* forward 5’-CCTGCGTGAGGACAACCCATAC-3’ and reverse 5’-GGTCTCATGCCAGCTGTTTCC-3’. *fhl2b* forward 5’-GCAAAAAGCCCATTGGCTGC-3’ and reverse 5’-CAGGTCTCATGCCAGCTGTTC-3’.

### BrdU Treatment

To label proliferating cell population in our model, zebrafish larvae were collected at 4dpf treated with BrdU at a concentration of 10mM and incubated overnight at 28.5 °C. At 5 dpf larvae were fixed in 4% PFA for 2 hours at room temperature. BrdU was counter-stained with anti BrdU-555 (1:500).

### Wound assay

3 dpf larvae were anesthetized in tricane in embryo medium and placed dorsal side up on a 2% agarose gel in a petri dish. Mechanical injuries were targeted to the somites on the dorsal side of embryos, above the cloaca, using a single stab with sharpened a glass capillary. This generated extensive muscle tissue damage localized to approximately 1 somite. The larvae were then washed in fresh embryo medium and reared until the appropriate stage. More than 95% of larvae survived this procedure.

### Imaging and automated measurements

Imaging of sectioned tissue and whole mount larvae was performed using a Nikon A1 confocal microscope (Nikon, Tokyo, Japan). Images are presented as five merged Z-stacks of representative areas or Z-depth in all figures. Automated measurements of EOM myofiber areas was performed using CellProfiler^65^.

### RNA-sequencing and analysis

For each group, 12 five months old and size matched (24 mm ± 3 mm) adult *desma^-/-^*;*desmb^-/-^*;*mylz2*:EGFP mutants and *WT^AB^;mylz2*:EGFP were euthanized by Tricaine. These were then swiftly subjected to EOM dissection, essentially as described^20^ but without paraformaldehyde fixation. All six EOMs still attached to the sclera were transferred into RNA-Later solution kept on ice and the EOMs were further cleaned, pooled and frozen in RNA-later. A piece of the lateral trunk including the slow domain myofibers from each fish was also excised and treated in the same manner. For EOM and trunk sequencing of 20 months old zebrafish, the same process was applied except the size matched animals were 31 mm ± 3 mm. In total, 144 fish were harvested. For 5 dpf zebrafish larvae, four groups of 10 larvae each containing sibling controls, sibling*; Tg(503unc:fhl2b-T2A-EGFP), dmd^-/-^* and *dmd^-/-^;Tg(503unc:fhl2b-T2A-EGFP)* larvae were cut diagonally distal to the swim bladder to exclude the majority of the gastrointestinal region and the head from the analysis. These larvae were then stored in RNA-later at -80°C until further use.

Pooled tissue was thawed and briefly rinsed in PBS before RNA extraction was performed using the TRIzol (Invitrogen, Thermo Fisher Scientific) reagent standard procedures and isolated in 15 µl of RNAse free H_2_O. Quality controls were performed using a bioanalyzer and a RIN value greater than 8 was considered acceptable for analysis. Library preparation was performed using the TruSeq Stranded mRNA Library Prep kit (cat#20020595, Illumina, Inc.), including poly-A selection, according to the manufacturer’s instructions. Unique index adapters were used (cat#20022371, Illumina, Inc. 15 cycles of amplification). The RNA-seq libraries were sequenced on a Novaseq 6000 Sequencing system (Illumina, Inc.) obtaining in average ∼78.5 million 150 paired end reads per library. Library preparation and sequencing was performed at SciLifeLab Stockholm.

Fastq files were quality controlled using FastQC (https://www.bioinformatics.babraham.ac.uk/projects/fastqc/) and raw reads were mapped to the zebrafish genome (GRCz11) using STAR (2.7.6a)^66^. Normalization and differential expression analysis were performed using DESeq2^67^, only genes with 10 or more reads were processed. Differentially expressed genes (DEGs, padj=<0.05) from different comparisons were intersected using the UpSet plot function from the ComplexHeatmap package^68^ and Gene Ontology (GO) analysis was performed using the clusterProfiler package^69^ (padj=<0.05) and plotted using the enrichR package. All statistical analysis related to RNA-sequencing data was performed within the R environment (version 4.2.3) using basic built-in functions and publicly available packages listed above. These are open source tools and can be accessed via Bioconductor (https://Bioconductor.org/).

### qPCR

Quantitative PCR was performed on using the same RNA-extraction method as described above on whole 5 dpf larvae. cDNA was synthesized using SuperScript IV (Invitrogen, Thermo Fisher Scientific). Primers used were forward 5’-GCTGCCAAGAATATCAGCGA-3’ and reverse 5’-TGCCTCCTCAGAGACTCATT-3’ for *desma*, forward 5’-ATGCAAGAGACCCAAGTCCA-3’ and reverse 5’-TCTTGCTCACAGCCTGGTTA-3’ for *desmb* and forward 5’-AAGGGGAACAGCTGGCATGA-3’ and reverse 5’-GTCGTGATAGGTCACGCCCC-3’ for *fhl2b*. β-actin was used as a reference gene using primers forward 5’-GCCTTCCTTCCTGGGTATGG-3’ and reverse 5’-CCAAGATGGAGCCACCGAT-3’.

The samples were run using an Applied Biosystems VIIA-7 Real Time PCR system (ThermoFisher Scientific) using FastStart universal SYBR green master mix (Roche).

### Physiological properties of zebrafish muscle force/tension relationship

Zebrafish larvae were examined with length-force experiments as previously described by Dou et al.^70^ and Li et. al.^71^. In brief, 5 and 6 dpf WT and *desma^-/-^*;*desmb^-/-^* larvae were euthanized and mounted with aluminum clips between a force transducer and a puller for length adjustment. The bath was perfused at 22°C in a MOPS buffered physiological solution with a pH of 7.4. The preparations were allowed to acclimatize in the solution for at least 10 min before initiating contractions using single twitch stimuli with 0.5 ms pulses at 2-min intervals and supramaximal voltage, via platinum electrodes placed on each side of the larvae. Measurements were initiated at slack length (Ls) and length was gradually increased by 10% steps every 4 min, between stimuli, until a decline in active force was observed. At each length, both active and passive force were recorded twice, and an average was calculated. The values were plotted against relative stretch (lambda=length/Ls).

### Spontaneous swimming and resistance swimming

5 dpf WT and *desma^-/-^*;*desmb^-/-^* zebrafish larvae were placed in a 48 well-plate inside the Viewpoint ZebraBox system (Viewpoint Behavior Technology) to determine spontaneous movement patterns. Larvae were carefully touched by the tail to ensure mobility and were subsequently allowed to acclimatize to the environment for 15 min before a 60 min protocol with a steady light was initiated. The movement thresholds were set to inactivity = 0 and large movements = 1. Inactivity, small movement and large movement counts, swimming distance and swimming duration were recorded.

For resistance swimming, 4 dpf larvae were placed in 1% methyl cellulose in 1x E3 medium and reared at 28.5°C over night. The following day, larvae were analyzed using fluorescent microscopy.

### Statistical analysis

Myofiber and myonuclei counts were performed in the whole slow domain of adult zebrafish trunks except when F310 positive myofibers were counted outside of the slow domain. F310 positive myofibers were then counted in 2x0.6 mm squares (20x magnification) just medial to the intermediate fast domain. The myofibers of each fish was counted twice, once for each side of the fish for all quantifications except for F310, where a total of four squares were counted, two for each side of the fish. The myofibers and myonuclei in the EOMs were counted separately in a total of two per fish. The medial rectus muscle was used. In embryonic experiments including Pax7, BrdU or TUNEL positive cells in the trunk muscle, somite number 13-22 was counted and the total number of positive cells were divided by the numbers of somites. For broken myofiber counts, somites 8-24 were counted. Number of larvae in the experiments are presented in the graphs.

All data was collected in Microsoft Excel and plotted in GraphPad Prism 10.0. Statistical analysis was performed using t-tests with Welch correction, p=<0.05 was considered significant (* p<0.05, ** p<0.005, *** p<0.0005, **** p<0.0001). All data are presented as mean ± standard error of mean (SEM). Kaplan-Meier log rank test was used to determine the difference between genotypes in the survival analysis. p=<0.05 was considered significant.

### Data availability

The RNA-seq datasets generated here have been deposited in Gene Expression Omnibus (GEO) under the accession number GSE242137.

## Legends

**Supplemental Figure 1.** A) Generation of *desma* and B) *desmb* CRISPR/Cas9 mediated knockout. C) Desmin immunolabeling in *desma^-/-^; desmb^-/-^* mutants and *desma^+/-^;desmb^+/-^* sibling control larvae at 3 dpf. D) *Tg(mylz2:EGFP)* positive fast and *Tg(smyhc1:tdTomato)* positive slow myofibers at 5 dpf in *desma^+/-^;desmb^+/-^*control and *desma^-/-^; desmb^-/-^* larvae cross section. E) Quantification of *Tg(mylz2:EGFP)* positive fast and *Tg(smyhc1:tdTomato)* positive slow myofibers at 5 dpf. F) Adult *desma^-/-^; desmb^-/-^*and WT (*desma^+/+^;desmb^+/+^*) controls at 5, 13 and 24 months. G) Cross-sections of EOMs in WT controls and *desma^-/-^; desmb^-/-^* 24 months old zebrafish immunolabeled for F310 (fast myofibers). H) Quantification of F310 positive myofibers relative to total myofiber population. I) Cross-sections of EOMs in WT controls and *desma^-/-^; desmb^-/-^* 24 months old zebrafish immunolabeled for S58 (slow myofibers). J) Quantification of S58 positive myofibers relative to total myofiber population. K) Cross-sections of WT and *desma^-/-^; desmb^-/-^*24 months old zebrafish trunk muscle immunolabeled for DAPI and Pax7. L) Quantification of Pax7/DAPI positive satellite cells (p=0.0008). M) Cross-sections of WT and *desma^-/-^; desmb^-/-^* 24 months old zebrafish trunk muscle immunolabeled for DAPI/PCNA. N) Quantification of PCNA/DAPI positive proliferating cells (p=0.0028). O) Cross-sections of WT and *desma^-/-^; desmb^-/-^* 24 months old zebrafish trunk fast domain muscle labeled for DAPI/TUNEL. Asterisk indicate myofibers with central nuclei. P) Quantification of myofibers with central nuclei in the fast domain of 24 months old zebrafish trunks showed significantly more regenerating myofibers in *desma^-/-^; desmb^-/-^* as compared to WT controls (p<0.0001). Dashed line in K) and M) indicates the myosepta separating the slow and fast muscle domain. Arrowheads in K), M) and O) indicate Pax7/DAPI, PCNA/DAPI and TUNEL/DAPI double positive nuclei, respectively. Open arrowheads in O) indicate TUNEL/DAPI positive nuclei on a different myofiber as compared to closed arrowheads. Data in violin plots (E, H, J, L, N, P) are presented as median (line) and quartiles (dashed line).

**Supplemental Figure 2.** A) Workflow for RNA-sequencing of 5 months and 20 months old WT (*desma^+/+^; desmb^+/+^*) and *desma^-/-^; desmb^-/-^* zebrafish larvae in *Tg(mylz2:EGFP)* background to visualize muscle tissue. B-I) Differential expression analysis performed for different pairwise comparisons: B) 5 months old EOMs in *desma^-/-^; desmb^-/-^ vs* WT, C) 5 months old trunk muscle in *desma^-/-^; desmb^-/-^ vs* WT, D) 20 months old EOMs in *desma^-/-^; desmb^-/-^ vs* WT, E) 20 months old trunk muscle in *desma^-/-^; desmb^-/-^ vs* WT, F) 5 months old EOMs *vs* trunk muscle in WT, G) 5 months old EOMs *vs* trunk muscle in *desma^-/-^; desmb^-/-^*, H) 20 months old EOMs *vs* trunk muscle in WT and I) 20 months old EOMs *vs* trunk muscle in *desma^-/-^; desmb^-/-^*zebrafish. Sample-to-simple distance heatmaps (left in B-I), PCA plots (middle in B-I) and MA plots (right in B-I) are displayed. MA plots show genes differentially expressed (DEGs, colored dots) for each comparison.

**Supplemental Figure 3.** Gene ontology terms enriched in the DEGs obtained from the comparisons A) 20 months old EOMs in *desma^-/-^; desmb^-/-^ vs* WT, B) 20 months old trunk muscle in *desma^-/-^; desmb^-/-^ vs* WT, C) 5 months old EOMs *vs* trunk muscle in WT, D) 5 months old EOMs *vs* trunk muscle in *desma^-/-^; desmb^-/-^*, E) 20 months old EOMs *vs* trunk muscle in WT and F) 20 months old EOMs *vs* trunk muscle in *desma^-/-^; desmb^-/-^*zebrafish. Yellow squares indicate myofiber related GO terms selected for further investigation for each comparison. CC: Cellular Compartment G) Fhl2 was found to be present mainly at the Z-disc in EOM myofibers studied using *Tg(mylz2:EGFP)* combined with Fhl2 antibodies and DAPI. Closed arrowheads indicate M-band positioning and open arrowheads indicate Z-disc positioning.

**Supplemental Figure 4.** Generation of A) *fhl2a* and B) *fhl2b* CRISPR/Cas9 mediated knockout zebrafish. Blue text shows CRISPR target site and red text shows deletions and faulty amino acids due to deletions. Asterisk indicates a premature stop codon. C) *in situ* hybridization of whole mount larvae at 5 dpf showing expression of *fhl2a* and D) *fhl2b.* E) Immunolabeling of Fhl2 in *desma^-/-^;desmb^-/-^;fhl2a^+/-^;fhl2b^+/-^*, *desma^-/-^;desmb^-/-^;fhl2a^-/-^*, *desma^-/-^ ;desmb^-/-^;fhl2b^-/-^* and *desma^-/-^;desmb^-/-^;fhl2a^-/-^; fhl2b^-/-^* larvae at 5 dpf, asterisk indicates lack of immunolabeling. Arrowheads in C-E) indicate EOMs, ventral view. F) *Tg(503unc:fhl2b-T2A-EGFP)* mosaic zebrafish larvae immunolabeled with Fhl2 antibodies. G-H) *Tg(503unc:fhl2b-T2A-EGFP)* mosaic zebrafish larvae with low and high mosaic expression labelled with phalloidin to visualize F-actin. Asterisk indicates somites with compromised integrity. I) Relative level of *fhl2b* in *Tg(503unc:fhl2b-T2A-EGFP)* zebrafish larvae with low and high stable EGFP expression. Data are presented as median (line) and quartiles (dashed line).

**Supplemental Figure 5.** A) Workflow for RNA-sequencing of 5 dpf sibling control (*dmd^+/-^*, *dmd^+/+^*), sibling control:*Tg(503unc:fhl2b-T2A-EGFP)*, *dmd^-/-^* and *dmd^-/-^:Tg(503unc:fhl2b-T2A-EGFP)* trunks. Larvae were cut distally to the swim bladder, indicated by the dashed line. B) Sample-to-sample distance heatmap and C) PCA plot for all four conditions. D-G) MA-plots showing differentially expressed genes (DEGs, green dots) for the comparisons D) *dmd^-/-^ vs* sibling control, E) *dmd^-/-^:Tg(503unc:fhl2b-T2A-EGFP) vs* sibling control, F) sibling control*:Tg(503unc:fhl2b-T2A-EGFP) vs* sibling control and G) *dmd^-/-^ vs dmd^-/-^ :Tg(503unc:fhl2b-T2A-EGFP)*.

**Supplemental Figure 6.** A-C) Gene Ontology terms enriched for the DEGs in the gene sets A, B and E (obtained by the intersection in Fig. 5A, H) are displayed in panels A, B and C, respectively. BP: Biological Process.**Supplemental Figure 7.** Examples from 5 sibling control, *dmd^-/-^* and *dmd^-/-^:Tg(503unc:fhl2b-T2A-EGFP)* larvae immunolabeled for A) Acetylated tubulin and B) α-bungarotoxin at 5 dpf, visualizing axons and NMJs, respectively.

**Supplemental Figure 8.** Sibling control (*dmd^+/+^*, *dmd^+/-^*), *dmd^-/-^*, *dmd^-/-^:Tg(503unc:EGFP)* and *dmd^-/-^:Tg(503unc:fhl2b-T2A-EGFP)* 5 dpf larvae immunolabeled for A) Pax7 showed B) a significantly decreased numbers of satellite cells in *dmd^-/-^:Tg(503unc:fhl2b-T2A-EGFP)* as compared to *dmd^-/-^* (p=0.0009) and *dmd^-/-^:Tg(503unc:EGFP)* (p=0.0018). C) Incorporation of BrdU (24h pulse) showed D) a significant decrease in proliferating cell numbers in *dmd^-/-^ :Tg(503unc:fhl2b-T2A-EGFP)* as compared to *dmd^-/-^*(p=0.0009) and *dmd^-/-^ :Tg(503unc:EGFP)* (p=0.0039) as well as E) decreased levels of proliferating satellite cells in *dmd^-/-^:Tg(503unc:fhl2b-T2A-EGFP)* as compared to *dmd^-/-^*(p=0.0017) and *dmd^-/-^ :Tg(503unc:EGFP)* (p=0.0035). F) TUNEL labeling showed a G) significant decrease in cell death in *dmd^-/-^:Tg(503unc:fhl2b-T2A-EGFP)* as compared to *dmd^-/-^* (p=0.0211) and *dmd^-/-^ :Tg(503unc:EGFP)* (p=0.0065). H) Immunolabeling of neutrophils using Mpx antibodies in uninjured 3 dpf sibling controls and *Tg(503unc:fhl2b-T2A-EGFP)*. I) Quantifications of DAPI/Mpx positive neutrophils in 3 dpf sibling controls and *Tg(503unc:fhl2b-T2A-EGFP)*. J) Immunolabeling of macrophages using Mfap4 antibodies in uninjured 3 dpf sibling controls and *Tg(503unc:fhl2b-T2A-EGFP)*. K) Quantifications of DAPI/Mfap4 positive macrophages in 3 dpf sibling controls and *Tg(503unc:fhl2b-T2A-EGFP)*. L) immunolabeling of satellite cells using Pax7 antibodies in uninjured 3 dpf sibling controls and *Tg(503unc:fhl2b-T2A-EGFP)*. M) Quantifications of DAPI/Pax7 positive satellite cells in 3 dpf sibling controls and *Tg(503unc:fhl2b-T2A-EGFP)*. N) Needle-stick injuries in 3 dpf sibling controls and *Tg(503unc:fhl2b-T2A-EGFP)* immunolabeled with Mpx antibodies from 2-18 hpi. O) Quantifications of DAPI/Mpx positive neutrophils in needle stick injured 3.08-3.75 dpf sibling controls and *Tg(503unc:fhl2b-T2A-EGFP)*. Data in O) is presented as mean ± SEM. Data in violin plots (B, D, E, G, I, K, M) are presented as median (line) and quartiles (dashed line). Avg. = average.

